# Antagonistic functions of CTL1 and SUH1 mediate cell wall assembly in *Arabidopsis*

**DOI:** 10.1101/2023.11.06.565903

**Authors:** Nguyen Thi Thuy, Hyun-Jung Kim, Suk-Whan Hong

**Author notes:** Corresponding author. E mail (S.W.H.).

## Abstract

Plant genomes contain numerous genes encoding chitinase-like (CTL) proteins, which have a similar protein structure to chitinase but lack the chitinolytic activity to cleave the *β*-1,4-glycosidic bond in chitins, polymers of *N*-acetylglucosamine. Mutations in *CTL1* in rice and *Arabidopsis* have been found to cause pleiotropic developmental defects, including altered cell wall composition and decreased abiotic stress tolerance, likely due to a reduction in cellulose content. In this study, we identified *suppressor of hot2-1* (*suh1*) as a genetic suppressor of the *ctl1^hot2-1^* mutation in *Arabidopsis*. The mutation in *SUH1* restored almost all *ctl1^hot2-1^* defects examined to nearly wild-type levels, with the exception of partial recovery of cellulose content in *ctl1^hot2-1^* mutants. *SUH1* encodes a Golgi-located type II membrane protein with glycosyltransferase (GT) activity, and its mutations lead to reduction in cellulose content and hypersensitivity to cellulose biosynthesis inhibitors, although to a lesser extent than *ctl1^hot2-1^* mutation. The *SUH1* promoter fused with the GUS reporter gene exhibited GUS activity in interfascicular fibers and xylem in stems, and this activity was significantly increased by the *ctl1^hot2-1^* mutation. Our findings provide genetic and molecular evidence that the antagonistic activities of CTL1 and SUH1 play an essential role in cell wall assembly in *Arabidopsis*.

**Highlight:** This study reports mutations in *suppressor of hot2-1* (*SUH1*) gene that rescues the defects caused by mutations in *CTL1*, which are characterized by stunted growth due to decreased cellulose levels.

## Introduction

Encompassing plant cells, cell walls (CWs) can be classified into two types, primary and secondary CWs, which differ in composition and function (Houston *et al*., 2016; Rui and Dinneny, 2019). The primary CW is relatively thin and flexible, consisting of cellulose microfibrils embedded in a matrix of various polysaccharides such as hemicelluloses and pectins, along with glycoproteins. It determines cell shape and controls cell expansion during growth. It also allows for cell-cell communication and contributes to the response to environmental stimuli. The secondary CW is thicker and more rigid than the primary CW. It is deposited on the inner side of the primary CW after cell expansion has ceased. The secondary CW is mainly composed of cellulose microfibrils, hemicelluloses, and lignin. Lignin, a polyphenolic compound, provides rigidity and waterproofing properties to the secondary CW. Consequently, the deposition of secondary cell walls in the interfascicular cells and xylem elements is crucial for providing structural support and facilitating water transport.

A wide range of mutations that affect cell wall synthesis have been documented to result in abnormal growth and development, which demonstrates the biological importance of the CW in plants. In *Arabidopsis*, isoxaben-resistant mutants (*ixr1* and *ixr2*) defective in cellulose synthesis of primary CWs exhibit severe defects in growth (Arioli *et al*., 1998; Scheible *et al*., 2001), while *irregular xylem* (*irx1* and *irx3*) mutations lead to defects in xylem development due to a decrease in cellulose content in secondary CWs (Chen *et al*., 2005; Taylor *et al*.,1999). In addition, the small stature of many mutants defective in non-cellulose polysaccharides such as pectin (Bouton *et al*., 2002; Liwanag *et al*., 2012) and hemicellulose (Persson *et al*., 2007; Xiao *et al*., 2016) provides genetic evidence of the physiological roles of the cell wall. The *Arabidopsis qua1* mutation, which causes a 25% decrease in pectin, results in dwarfism and reduced cell adhesion (Bouton *et al*., 2002). The *Arabidopsis irx8* mutant plants with a significant decrease in hemicellulose, such as homogalacturonan and xylan, display dwarfism and a reduction in secondary cell wall thickness (Persson *et al*., 2007). Moreover, rice *brittle culm* (*bc*) mutants with reduced mechanical strength of internodes also demonstrate the essential role of cell walls in growth and development (Zhang and Zhou, 2011).

Chitinases comprise a subfamily of glycosidic hydrolases (GHs) that catalyze the hydrolysis of β-1,4 glycosidic bonds in amino polysaccharides such as chitin and chitooligosaccharides (Kesari *et al*., 2015). They are well-known for their involvement in plant defense, as they release chitin fragments from fungal cell walls, which can subsequently stimulate plant innate immune responses. However, recent studies have revealed that chitinase-like (CTL) proteins, which have a similar structure to chitinase but lack chitinolytic activity, participate in CW assembly and affect plant growth and development. Mutations in *CTL1* in *Arabidopsis* result in reduced root length (Hauser *et al*., 1995), anion-altered root morphology (Hermans *et al*., 2010), ectopic deposition of lignin, abnormal cell shapes with incomplete cell walls in the pith (Zhong *et al*., 2002), and reduced tolerance to abiotic stresses such as high temperature and salinity (Hong *et al*., 2003; Kwon *et al.,* 2007). Sánchez-Rodríguez *et al*. (2012) demonstrated that AtCTL1 is secreted from cells through the secretory pathway and has the ability to bind glucan-based polymers *in vitro*. Based on these findings, they postulate that apoplastic CTL1 interacts with cellulose microfibrils and hemicelluloses, influencing cell wall assembly in *Arabidopsis*. Interestingly, a mutation in *BC15*/*OsCTL1*, a rice ortholog of *Arabidopsis CTL1*, was also found to decrease cellulose content in rice (Wu *et al*., 2012). These findings suggest that CTL1 activity in rice also participates in cell wall assembly, similar to *Arabidopsis*. However, little is known about the molecular mechanisms by which *CTL1* mediates cell wall synthesis in *Arabidopsis* and rice.

In this report, we employed genetic approaches to further investigate the role of *CTL1* in cell wall assembly, which in turn affects growth and development in *Arabidopsis*. We identified *suppressor of hot2-1* (*suh1*), which ameliorates the short hypocotyls of etiolated *ctl1^hot2-1^* seedlings. We report that *SUH1* encodes a Golgi-localized type II membrane protein with glycosyltransferase activity, potentially involved in glycan synthesis. Our findings provide novel insights into the function of *CTL1* in cell wall synthesis.

## Materials and Methods

Unless stated otherwise, all mutants and transgenic lines used in this study were obtained from the *Arabidopsis* Columbia-0 ecotype (Col-0). We obtained the *Arabidopsis thaliana* SAIL_912_D02 T-DNA insertion line (*suh1-4*) with T-DNA inserted in the first intron of *At5g14550* from the *Arabidopsis* Information Resource (TAIR). Genomic DNA was extracted from individual plants. To confirm T-DNA insertion, PCR amplification was performed using gene-specific primers and the T-DNA left border primer LBb1 (Supplementary Table S*2*). The precise position of the T-DNA insertion was determined by sequencing the PCR products with the T-DNA left border primer LBb1.

### EMS Mutagenesis and Isolation of the Suppressor of *ctl1^hot2-1^* Mutation

Approximately, 10,000 *ctl1^hot2-1^* seeds (M_1_ seeds) were imbibed overnight in water at 4 °C and then incubated for 12 h in 50 mL of 0.25% ethyl methane sulfonate (EMS). The EMS was subsequently washed several times with water. The M_2_ seeds were independently harvested from 100 trays, each containing about 100 M_1_ plants. Because dark-grown *ctl1^hot2-1^* seedlings display a short hypocotyl, about 120,000 M_2_ seeds were grown in dark to identify three independent seedlings with a long hypocotyl. Then, it was confirmed that these three dark-grown suppressors also showed a long hypocotyl in the successive M_3_ generation, and they were designated as *suh* (<u>su</u>ppressor of *<u>h</u>ot2*) mutants. In order to determine the inheritance characteristics of the suppressor mutation, three *suh* mutant plants were crossed with the parental *ctl1^hot2-1^* plants, and all pairwise combinations of the three *suh* mutants were crossed to conduct allelism tests.

### Map-based Cloning of *SUH1* Gene

To perform map-based cloning of the *suh1* mutation, we looked for a *ctl1* mutant allele in *Landsberg erecta (*Ler) ecotype by conducting a genetic screening of EMS-mutagenized Ler seeds. Complementation test and sequencing analysis revealed that a new *ctl1* allele (Ler) harbors the same mutation as the *ctl1^hot2-1^* (G881A).

To generate a mapping population, *ctl1^hot2-1^ suh1-1* mutant (Col-0) was crossed with Ler-derived *ctl1* plants named *ctl1^hot2-3^*. The resulting F_1_ plants were self-pollinated, and DNA was extracted from individual F_2_ plants displaying the wild-type phenotype of seedlings grown in dark. PCR was performed using simple sequence length polymorphism (SSLP) markers (Bell and Ecker,1994). The recombination frequency between the *SUH1* locus and the SSLP makers was used to determine the genetic map position of the *SUH1* locus.

### Semi-Quantitative RT-PCR and cDNA Synthesis

Total RNA was extracted from stems of *Arabidopsis* and rice using the Ribospin^TM^ kit from GeneAll (https://geneall.com/). For RT-PCR, 2–5 μg of total RNA was reverse transcribed using a Superscript First-Strand Synthesis system (Invitrogen, Carlsbad, CA, USA). The quantitative RT-PCR was performed using the *suh1-4* RT primers (Supplementary Table S2) and *Arabidopsis actin2* as an endogenous reference. PCR products were visualized by agarose gel electrophoresis with EtBr.

First-strand cDNA was synthesized from 5 μg of total RNA using the PrimeScript^TM^II cDNA Synthesis kit from Takara (https://takara.com/). The full-length cDNAs of *SUH1* and *BC10* were amplified using the Taq LA DNA polymerase PCR kit from Takara (https://takara.com/) with cDNA-specific primers (Supplementary Table S2). The resulting fragments were cloned into the pGEM-T easy vector and sequenced using an ABI 3730 automated sequencer (Applied Biosystems).

### Construction of Various Vectors and Plant Transformation

The 1,060-bp promoter fragment, spanning from −1,060 to −1 of the *SUH1* genomic DNA, was amplified (the A site of ATG, the translation start codon, was designated as +1). The *pSUH1::GUS* construct was generated by replacing the 35S promoter of the binary vector pBI121 with a fragment containing the *SUH1* promoter. To generate the constructs used for complementation tests, the full-length cDNAs of *SUH1* and *BC10* were placed under the control of *SUH1* promoter sequence (1,060 bp) in the binary vector pBI121. The resulting binary vectors (*pSUH1::SUH1* and *pSUH1::OsBC10*) were introduced into the *Agrobacterium tumefaciens* strain GV3101 for *Arabidopsis* transformation using the floral dip method (Clough and Bent, 1998). Seedlings from the T_1_ generation were selected on half-strength MS medium containing 30 μg/mL kanamycin. T_3_ progenies homozygous for each transgene were isolated and three independent lines for each construct were selected for further examinations.

### Cell Wall Composition Assay and Histology

The cellulose content of inflorescence stems (5-cm-long, bottom) was determined in 6-week-old wild-type and mutant plants using the Updegraff assay (Kumar and Turner, 2015). Samples were incubated in 70% ethanol at 70 °C for 1 h, washed with 100% acetone, and dried overnight. After adding the acetic nitric agent (8:1:2, acetic acid: nitric acid: water), the samples were placed in boiling water bath for 30 min, precipitated by centrifugation at 14,000 rpm for 15 min, and incubated in 67% sulfuric acid in a boiling water bath for 5 min. Finally, the anthrone-sulfuric acid colorimetric assay was performed to analyze the samples. The absorbance was measured at 620 nm using a spectrometer (UV-1600, Shimadzu). The cellulose contents were expressed as the percentage of cellulose in the cell wall composition, with glucose standards.

To stain lignin, pectin, and arabinogalactan proteins, hand-cut sections of inflorescence stems were incubated in 1% (w/v) phloroglucinol in 18% HCl, 0.01% (w/w) ruthenium red or 2 mg/mL β-glycosyl Yariv agent in 0.1 M NaCl, respectively. After staining, the samples were washed with deionized water, and observed under a Carl Zeiss AX10 microscope (Zeiss Corp, Germany).

To determine the shapes of the pith cells, the inflorescence stems of 6-week-old wild-type and mutant plants were fixed in 2.5% glutaraldehyde at 4 °C overnight. Tissues were gradually dehydrated in ethanol and embedded in Historesin (Leica Microsystems, France) according to the manufacturer’s instructions. Transverse sections (5-μm-thick) were prepared using an ultramicrotome (LEICA, Germany), stained with 1% toluidine blue, and examined under a Carl Zeiss AX10 microscope (Zeiss Corp, Germany).

### Treatment with Cellulose Biosynthesis Inhibitors and Abiotic Stress

To determine their sensitivity to cellulose biosynthesis inhibitors (CBIs), the sterilized *Arabidopsis* seeds were grown vertically on half-strength MS medium supplemented with the indicated concentrations of 2,6-dichlorobenzonitrile (DCB) or isoxaben (ISX) in 0.5% dimethyl sulfoxide (DMSO). An equal volume of DMSO was used as a control. After a growth period of 10 days, the primary root growth was measured.

To assess thermotolerance, seedlings were grown vertically for 2.5 days in dark, subjected to heat-stress for 2 h at 45 °C, further grown for 2.5 more days in dark, and analyzed. To apply salinity stress, 3-day-old seedlings grown vertically on half-strength MS medium were transferred to a medium with or without the indicated concentrations of NaCl. After 7 *days* of *growth* under long-day conditions (16-h light/8-h dark), the primary root lengths of wild-type and mutant plants were measured.

### Histochemical Assay for GUS Activity

Histochemical analysis of β-glucuronidase (GUS) activity was conducted as described previously (Jefferson *et al*., 1987). GUS enzyme activity in transgenic plants was determined by staining with 1 mg/mL X-Gluc (Duchefa, the Netherlands) as the substrate. The GUS-stained tissues shown in this report represent the typical results obtained in three independent transgenic lines.

### Assay of C2GnT Enzyme Activity

The C2GnT assay was performed as described previously (Bierhuizen and FuKuda M, 1992). To this end, *SUH1* and *BC10* were fused in-frame with *EGFP* using the pEGFP-N1 vector. The plasmids were transiently transfected into Chinese hamster ovary (CHO) cells. After 48 h, the cells were washed with phosphate-buffered saline, and suspended in lysis buffer (10 mM TrisCl at pH 8.0, 0.1 mM EDTA, 5 mM DTT, 0.9% NaCl, and 1% Triton X-100). The cell lysate was centrifuged at 1000 *g* at 4 °C for 10 min. The supernatant was then stored in aliquots of 250 μL at –80 °C until further use. Protein concentration was determined using a Bio-Rad protein assay with bovine serum albumin (BSA) as the standard. As the accepter, Galβ1 → 3GalNAcα1 → p-nitrophenyl (Sigma) was used for the C2GnT assay. Non-transfected and empty pEGFP-N1-transfected CHO cells were used as the negative control group. The reaction mixtures contained 50 mM HEPES-NaOH at pH 7.0, 1 mM UDP-[glucosamine-U-^14^C] GlcNAc (3.7 kBq, Amersham Pharmacia Biotech), 1 mM acceptor oligosaccharide, 0.1 M GlcNAc, and 5 mM DTT. After incubation for 3 h at 37 °C, the reaction products were adjusted to 0.25 M ammonium formate at pH 4.0, and applied to a C18 reverse phase column (Alltech Associates Inc, IL, United States). After washing the column with the same solution, the product was eluted with 70% methanol. The radioactivity was measured using scintillation counting.

### Tobacco Transient Expression and Confocal Microscopic Analysis of SUH1–GFP Fusion Proteins

To investigate its subcellular location, *SUH1* was fused in-frame with the GFP-encoding sequence of the pCAMBIA1300 vector. The ER marker mCherry-HDEL and the Golgi marker MAN49-mCherry were obtained from the Daegu Gyeongbuk Institute of Science and Technology (DGST). Transient transformations of *Nicotiana benthamiana* with *A. tumefaciens* strain (GV3101) containing the *SUH1-GFP* construct and organelle markers were conducted. The *Agrobacterium* cells were incubated in an infiltration buffer (10 mM MgCl_2_, 10 mM MES at pH 5.9, and 150 μM acetosyringone) for 3 h at room temperature. Before infiltration, the bacterial suspension was mixed with an equal volume of a bacterial suspension harboring pBin61-P19 to co-introduce the RNA-silencing suppressor gene into the cells. The mixture of *Agrobacterium* suspensions was then infiltrated using a 5-mL syringe without a needle into the abaxial side of the second, third, and fourth leaves of 6-week-old *N. benthamiana* plants grown in a growth chamber at 25 °C under long-day conditions (16-h light/8-h dark). The agroinfiltrated plants were placed back into the same growth chamber and examined for fluorescence expression using a Carl Zeiss LSM 710 confocal scanning microscope.

## Results

### Mutations in *SUH1* restore the growth retardation of dark-grown *ctl1^hot2-1^*seedlings

In order to investigate the molecular mechanisms underlying *CTL1*-mediated cell wall assembly in *Arabidopsis*, a genetic screen was conducted on over 120,000 M_2_ seedlings of ethyl methanesulfonate (EMS)-mutagenized *ctl1^hot2-1^* plants. The aim was to identify secondary mutations that could restore the short hypocotyl of 5-day-old dark-grown *ctl1^hot2-1^* seedlings. Three independent suppressors were isolated based on their ability to elongate their hypocotyl to wild-type length in successive generations (Fig. 1A, B). In addition, these suppressors also restored the abnormal morphology of light-grown *ctl1^hot2-1^* mutants, including the presence of numerous lateral branches and shorter stature of the aerial parts, to that of wild-type plants (Fig. 1C). Pair-wise crosses of three suppressors demonstrated that the three mutations affected a single gene, leading to their designation as *suh1-1*, *suh1-2*, and *suh1-3* (Supplementary Table S1). Their crosses with *ctl1^hot2-1^*mutants also revealed that the restoration of hypocotyl elongation is inherited in a recessive manner.

**Figure 1.**
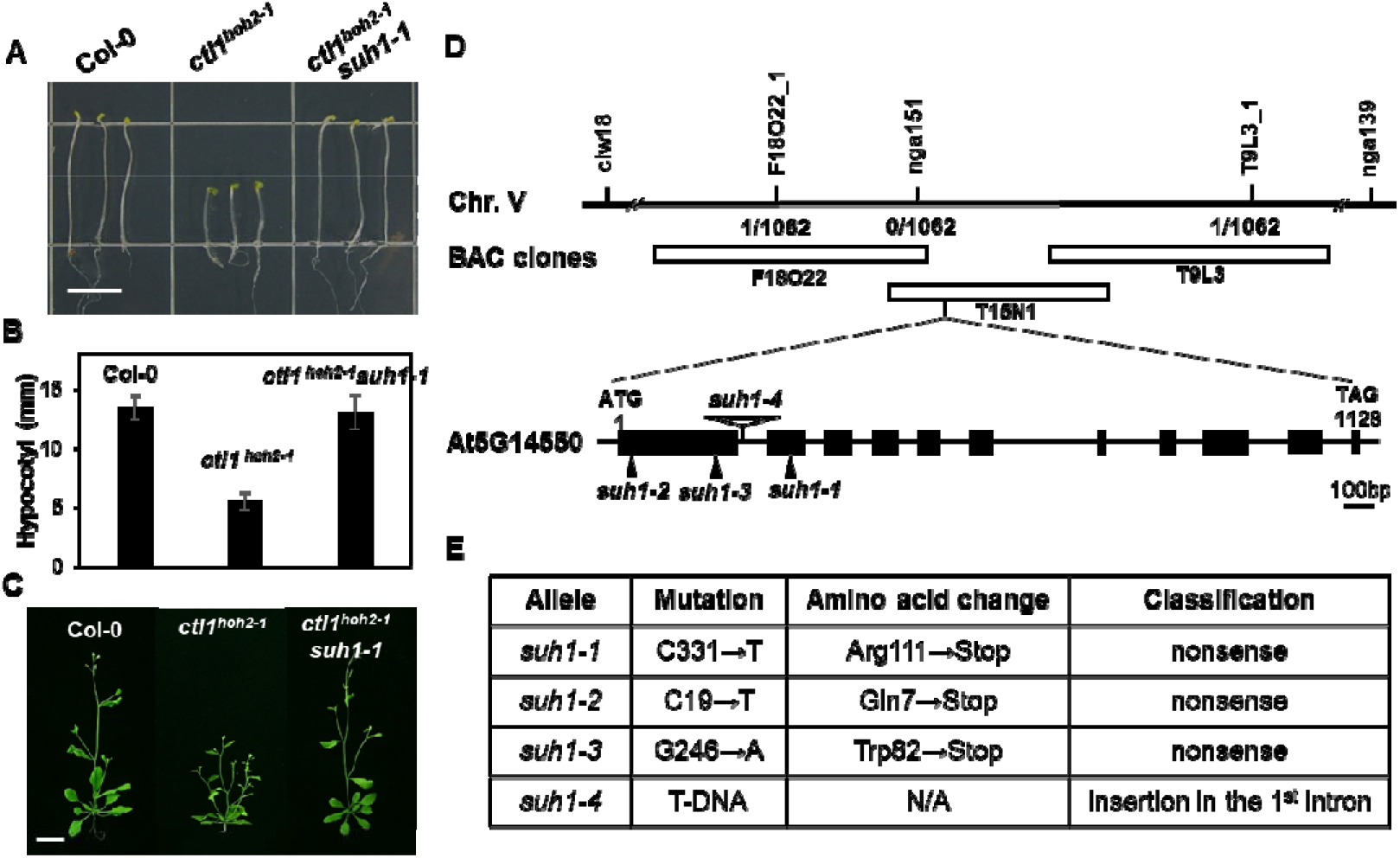
Mutations in *SUH1* restore growth defects of *ctl1^hot2-1^* mutants. (A) Representative five-day-old dark-grown seedlings of Col-0, *ctl1^hot2-1^*, and *ctl1^hot2-1^ suh1-1* plants. Scale bar, 5 mm. (B) Quantification of hypocotyl length in etiolated seedlings shown in panel *A*. Data are expressed as the mean ± standard error (*SE)* of three replicates. (C) Representative six-week-old light-grown Col-0, *ctl1^hot2-1^*, and *ctl1^hot2-1^ suh1-1* plants. Scale bar, 1 cm. (D) Map-based cloning of *SUH1*. *SUH1* was mapped to a region on chromosome 5 between two simple sequence length polymorphism (SSLP) markers, nga151 and T15N1_1, which is covered by three overlapping BAC clones: F18O22, T15N1, and T9L3. The numbers of recombinants obtained from 536 F_2_ individual plants (1,062 chromosomes) are indicated under each marker. The genomic structure of *SUH1* is shown with black boxes and lines indicating exons and introns, respectively. Point mutations are represented by solid arrowheads; T-DNA insertion by an open arrowhead. (E) Descriptions and predicted molecular effects of the three *suh1* point mutations and of the T-DNA insertion. N/A indicates not applicable.

### Map-based cloning of *SUH1*

The *SUH1* locus was mapped to a 53-kb segment on chromosome 5 (Fig. 1D). Through sequencing analysis of *suh1-1* mutants, we identified a premature stop codon caused by a substitution of Arg at position 111 in the second exon of *At5g14550*, which consists of 11 exons and 10 introns (Fig. 1E). Nonsense mutations were also detected in *suh1-2* and *suh1-*3, where Gln^7^ and Trp^82^ in the first exon of *At5g14550* were replaced with stop codons, respectively (Fig. 1D, E). In addition, we identified a fourth mutant allele, *suh1-4*, which carried T-DNA in the first intron of *At5g14550* (Fig. 1D, E). No *SUH1* transcripts were detected in *suh1-4* mutant plants (Supplementary Fig. S1A), and *suh1-4* was confirmed to restore the growth defects of *ctl1^hot2-1^* mutants under both dark and light conditions. The introduction of *SUH1* cDNA driven by its own promoter in *ctl1^hot2-1^ suh1-1* mutants resulted in a reversion to the *ctl1^hot2-1^* phenotype under both dark and light conditions (Supplementary Fig. S1B, C), providing further evidence supporting that the mutation in *SUH1* (*At5g14550*) is responsible for the suppression of the *ctl1^hot2-1^* phenotype.

Hématy *et al*. (2007) showed that mutations in *THE1*, which encodes a receptor-like kinase, partially restore not only the short hypocotyl of etiolated *cesa6^prc1-1^* carrying a null mutation in *CESA6* (resulting in reduced cellulose levels) but also that of *pom1-2* seedlings, which is another allele of *CTL1* in *Arabidopsis*. To examine whether *suh1* can also restore the short hypocotyl of etiolated *cesa6^prc1-1^* mutant, *suh1-4 cesa6^prc1-1^* double mutants were generated by crossing *suh1-4* and *cesa6^prc1-1^*mutants. When grown in dark, the appearance of *suh1-4 cesa6^prc1-1^*mutant seedlings was indistinguishable from that of *cesa6^prc1-1^*mutants (Supplementary Fig. S2), suggesting that *suh1-4* is unable to rescue the growth defects of *cesa6^prc1-1^*mutants. This implies that *SUH1* is involved in cell wall assembly in association with *CTL1*, rather than directly participating in cellulose synthesis in *Arabidopsis*.

### Mutations of *SUH1* restore multiple defects caused by *ctl1^hot2-1^*to nearly wild-type levels

To examine the effect of *suh1* on the various defects linked to *CTL1* mutation, we compared the phenotypes of plants carrying four combinations of *ctl1^hot2-1^*and *suh1-4*. First, we confirmed that *suh1-4* restored the root growth of *ctl1^hot2-1^*mutants to wild-type levels (Fig. 2A). As shown in Figure 2B, *suh1-4* suppressed the increase in root hair density in *ctl1^hot2-1^* plants; the root hair density of *ctl1^hot2-1^ suh1-4* plants was indistinguishable from that of wild-type and *suh1-4* plants. Next, we examined whether *suh1* could restore changes in cell wall composition and abnormal cell shape in *ctl1^hot2-1^*mutants. Using the Updegraff assay (Kumar and Turner, 2015), we found that the cellulose content in *ctl1^hot2-1^* mutants decreased to 66.9% of that in wild-type plants and that *suh1* restored the cellulose content in *ctl1^hot2-1^*mutants to 81.6% of that in wild-type plants (Fig. 2C). These results indicate that cellulose biosynthesis in *ctl1^hot2-1^* mutants is partially restored by *suh1-4*. Interestingly, *suh1-4* mutants also exhibited a statistically significant decrease in cellulose content compared with wild-type plants, despite being indistinguishable from wild-type plants in growth and development under non-stress growth conditions (Fig. 2A and supplementary Fig. S2B). These findings suggest that the *suh1* single mutation does not affect growth but may have non-negligible effects on cellulose biosynthesis under optimal growth conditions. In addition to the partial restoration of cellulose content, histochemical staining of cell wall elements revealed that the deposition patterns of lignin (Fig. 2D), pectin (Fig. 2E), and arabinogalactan proteins (AGPs; Fig. 2F) in inflorescence stems of *ctl1^hot2-1^ suh1-4* mutants were indistinguishable from those in *suh1-4* and wild-type plants, unlike *ctl1^hot2-1^*plants, which displayed ectopic depositions. Cross-sections of mature stems showed that *suh1-4* completely rescued the larger and irregularly shaped cells in the pith of *ctl1^hot2-1^* mutants to a wild-type phenotype (Fig. 2G).

**Figure 2.**
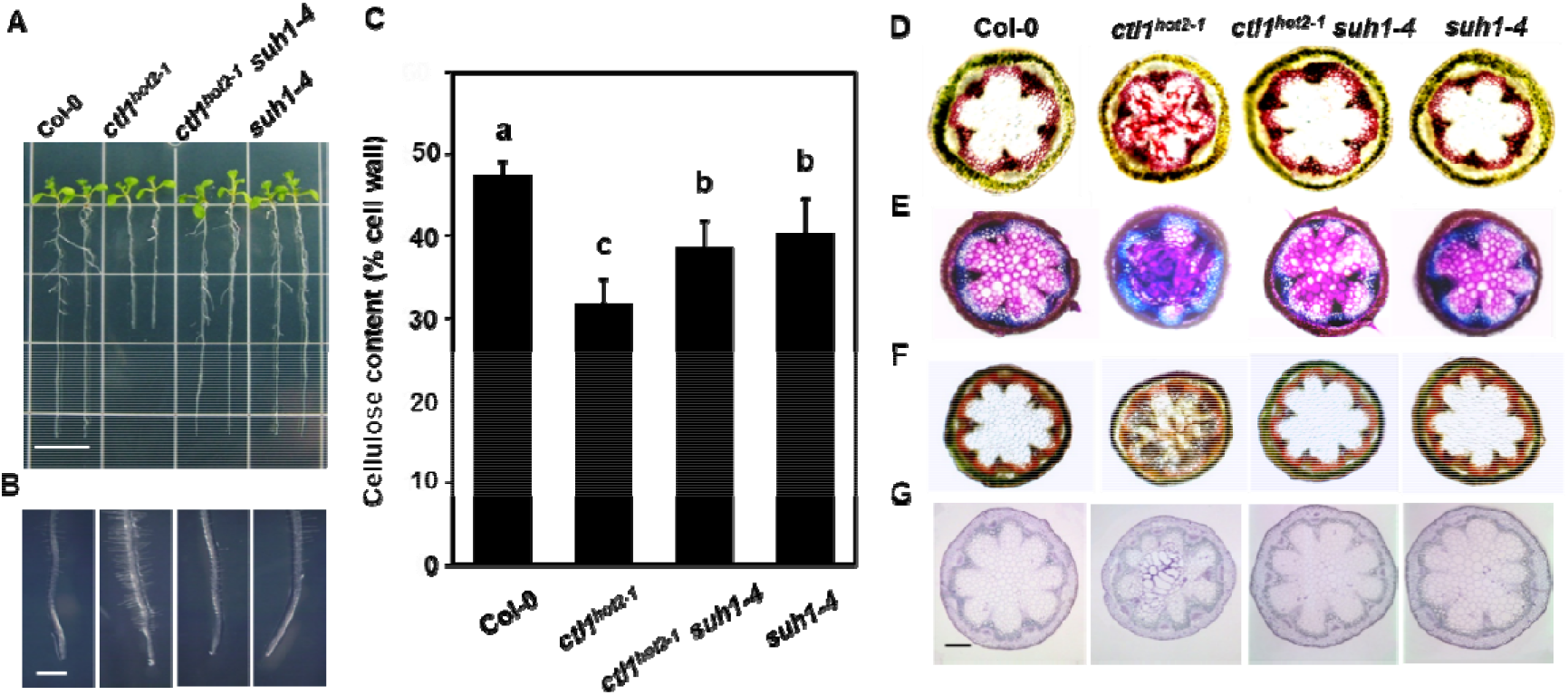
The defects in *ctl1^hot2-1^*mutants are restored by the *suh1* mutation. Representative images of the primary root (A) and root hair (B) of 10-day-old wild-type and mutant plants grown vertically on half-strength MS medium. Scale bar, 1 cm (A) and 1 mm (B). (C) Total cellulose contents (% cell wall) in inflorescence stems of 6-week-old plants. Data are expressed as the mean ± *SE* (11 samples). Statistically significant differences are indicated with different letters (one-way ANOVA followed by Tukey’s test, P < 0.05). Hand-cut sections of stems of 6-week-old wild-type and mutant plants stained with phloroglucinol-HCl (D), toluidine blue (E), or β-glucosyl Yariv (β-GlcY) reagent (F). Phloroglucinol-HCl stains lignin in red. Toluidine blue stains the cell wall containing lignin and pectin in blue-green and pink, respectively. β-GlcY stains arabinogalactan proteins (AGPs) in reddish brown. (G) Transverse sections of inflorescence stems of 6-week-old wild-type and mutant plants. Sections were prepared at the first internode of stem. Scale bar, 200 µm (D)-(F).

Moreover, *suh1-4* also restored the reduced tolerance to high temperature (Supplementary Fig. S3A, B) and salinity stress (Supplementary Fig. S3C, D) of *ctl1^hot2-1^* plants. These results indicate that *suh1-4* can effectively restore almost all defects caused by *ctl1^hot2-1^* to nearly wild-type levels, with the exception of partial recovery of cellulose content in *ctl1^hot2-1^*. These findings suggest that there may be a critical threshold for cell wall integrity, below which both growth and abiotic stress tolerance are affected in *Arabidopsis*. While *ctl1^hot2-1^*appears to affect cell wall integrity to levels below this threshold, resulting in growth defects, *suh1* is likely to have minimal impact on growth and development, possibly due to the maintenance of cell wall integrity above the threshold.

### The response of cellulose biosynthesis inhibitors is altered by *suh1*

The discovery that *suh1-4* affects cellulose synthesis prompted us to investigate the response of wild-type and mutant plants to chemicals that inhibit cellulose synthesis. To this end, we examined root growth in 10-day-old light-grown seedlings on half-strength MS media supplemented with cellulose biosynthesis inhibitors (CBIs) such as 2,6-dichlorobenzonitrile (DCB) and isoxaben (ISX). The primary root growth in wild-type plants on medium with the lowest concentration of CBIs was more than 70% of that in control plants on medium without CBIs, while the primary roots of *ctl1^hot2-1^* mutants barely grew under the same conditions (Fig. 3). As expected, the primary roots of *ctl1^hot2-1^ suh1-4* mutants exhibited significant elongation compared to those of *ctl1^hot2-1^* mutants at all inhibitor concentrations tested, suggesting that *suh1-4* partially restores root growth retardation in *ctl1^hot2-1^* mutants on MS media containing CBIs. Furthermore, *suh1-4* mostly exhibited root growth comparable to that of *ctl1^hot2-1^ suh1-4* at all concentrations of CBIs, indicating that the upper limit of the response of *ctl1^hot2-1^ suh1-4* is determined by *suh1-4*. In addition, *suh1-4* was more sensitive to DCB than to ISX. ISX is known to inhibit cellulose synthase, while DCB affects microtubule-associated protein (MAPs) that play a key role in vesicle transport (Wormit *et al*., 2012). Therefore, these results suggest that *suh1* may be more sensitive to disruption of vesicle transport rather than directly affecting cellulose synthesis.

**Figure 3.**
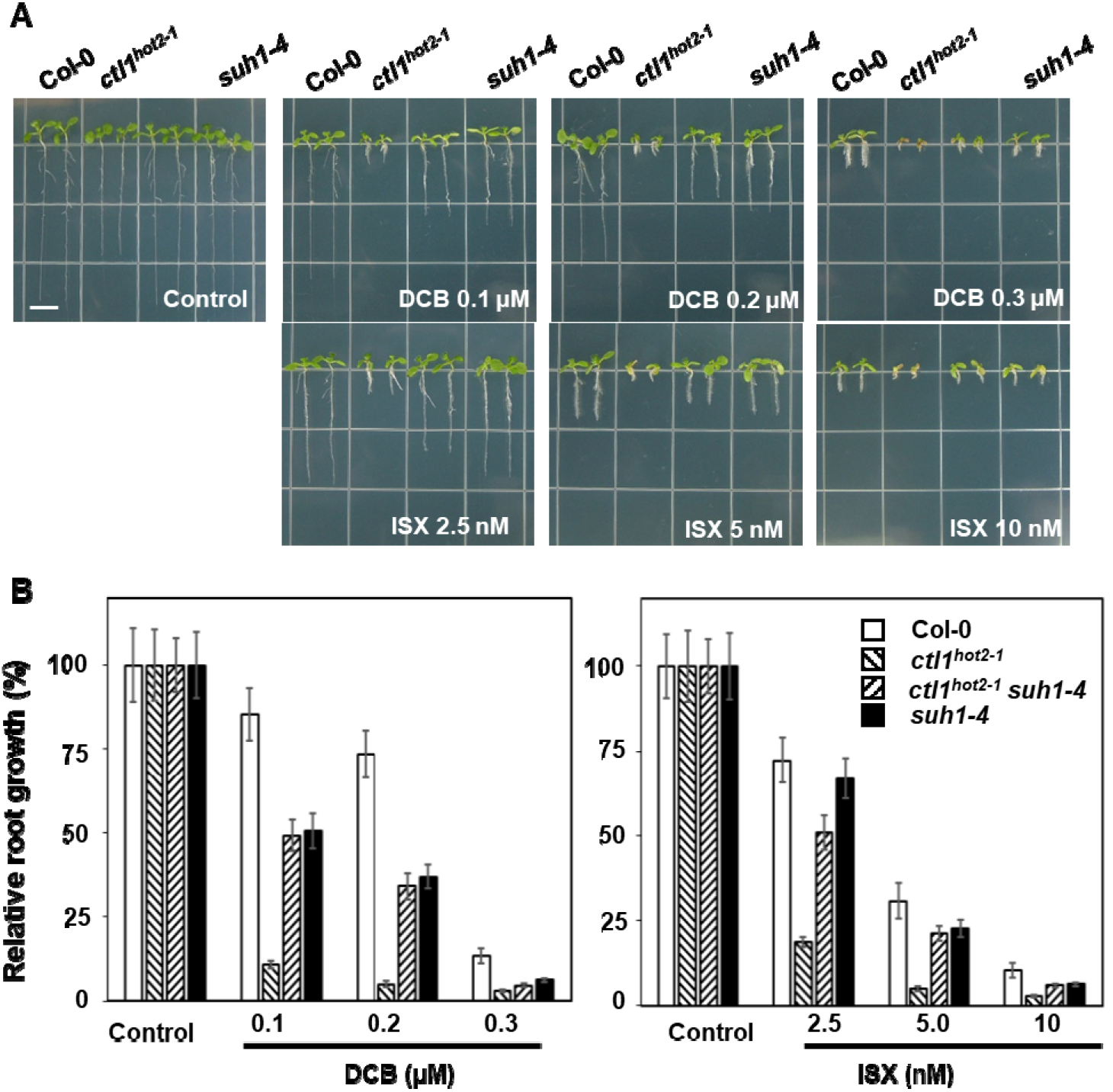
Responses of wild-type and mutant seedlings to CBI treatment. Upon CBI treatment, the upper limit of root growth in *ctl1^hot2-1^ suh1-4* plants is determined by the *suh1-4* mutation. (A) Representative 10-day-old seedlings of wild-type and mutant plants on half-strength media containing different concentrations of DCB or ISX. Scale bar, 0.5 cm. (B) Quantification of root lengths of the seedlings described in (A). Data are presented as the means ± SE of three replicates of 15 seedlings each.

### *SUH1* encodes a Golgi-localized type II membrane protein containing the domain of unknown function 266

The *SUH1* cDNA has an open reading frame of 1134 nucleotides, which encodes a polypeptide of 377 amino acids with a predicted molecular mass of 44551.5 Da. Sequence analyses using Pfam (Finn *et al*., 2006) revealed an N-terminal transmembrane domain (TM, amino acids 20–39) followed by a C-terminus containing the domain of unknown function 266 (DUF266, amino acids 66–322). The multiple alignment of various DUF266-containing proteins showed that SUH1 is most similar to the rice BC10 (Fig. 4A). Bioinformatic analyses suggest that DUF266-containing proteins have structural similarities to the Leukocyte core-2 β1, 6N-acetylglucosaminyltransferase (C2GnT-L; Fig. 4B), which is a member of the glycosyltransferase 14 (GT14) family (Hansen *et al*., 2009). DUF266-containing proteins are structurally related to the GT14 family and have been found in almost all plants (Ye *et al*., 2011). However, the study of *Brittle Culm 10* (*OsBC10*), which encodes a type II membrane protein containing DUF266 in rice, was the first report on their functional characterization (Zhou *et al*., 2009). The catalytic region of human C2GnT contains three functional domains (Hansen *et al*., 2009; Fig. 4B). The first domain is the Rossmann-type nucleotide-binding domain (amino acids 125–225 in C2GnT) followed by the substrate-interaction domain (amino acids 286–345 in C2GnT) and the final domain that binds to the diphosphate group of the nucleotide (amino acids 396–424 in C2GnT). We identified counterparts exhibiting high similarity to the three functional domains of C2GnT in DUF266 proteins (Fig. 4B). In particular, it is interesting to note that the Glu^320^ and Lys^401^ residues of C2GnT, which are involved in catalysis and nucleotide binding, are also identified in DUF266 proteins. These residues are indicated by an asterisk and a circle, respectively (Fig. 4B).

**Figure 4.**
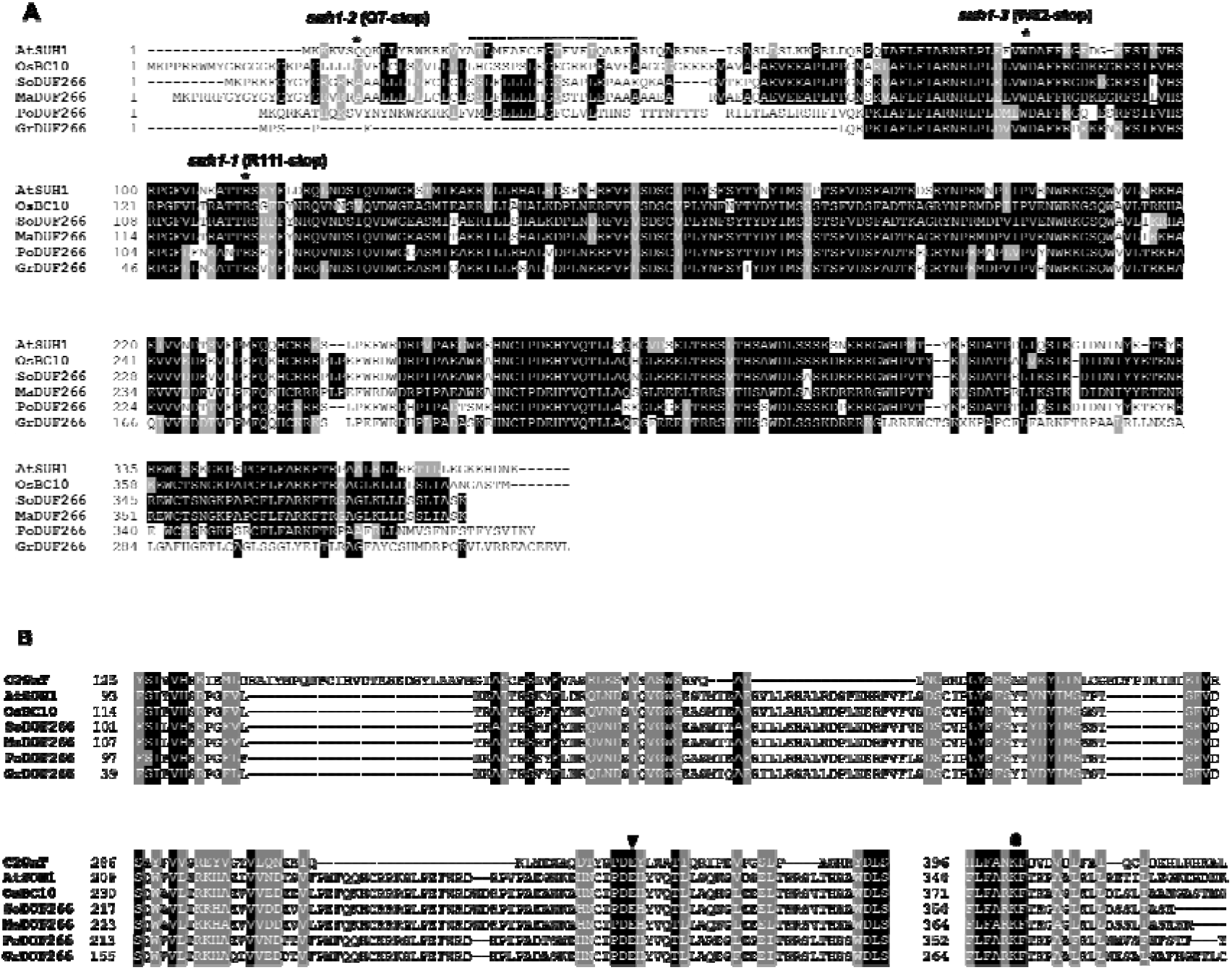
*SUH1* encodes a transmembrane protein containing a domain of unknown function 266 (DUF266) with high similarity to the GT14 protein. (A) Multiple alignment analysis of SUH1 and predicted DUF266-containing proteins from plants. The predicted amino acid sequences from *Arabidopsis* (SUH1, B6IDH4), rice (BC10, Q65XS5), sorghum (So, C5Z134), maize (Ma, B4FL81), poplar (Pp, B9GHR6), and grape (Gr, A5AN07) were aligned using the ClustalW program (mobyle.pasteur.fr/cgi-bin/portal.py). White letters on a black background indicate residues that are invariant, and other conserved amino acids are shaded in gray. The predicted transmembrane region (amino acids 20–39) is indicated by a dotted line, and the positions of *suh1* point mutations are indicated by asterisks. (B) Partial amino acid sequence alignment of DUF266 protein members and the human Core2 1,6 β-*N*-acetylglucosaminyltransferase (Q02742) of the GT14 family. Bioinformatic studies revealed structural similarities and invariant amino acid residues conserved between DUF266 proteins and leukocyte type core-2 β-1,6-*N*-acetylglucosaminyltransferase (C2GnT-L), a member of the GT14 family, suggesting a distant relationship between DUF266 proteins and GT14. The first region presumably corresponds to the Rossmann-fold motif for nucleotide binding. The second region is a structural domain that interacts with both the donor and acceptor substrates. Of particular interest is the conservation of the putative catalytic residue Glu-320 of C2GnT in DUF266 members. The putative catalytic residue (Glu-320 in C2GnT) is marked with an inverted triangle. The third region at the C-terminus of the catalytic domain shows three invariant residues. One of them, Lys-401 in C2GnT (marked with a circle), which is expected to interact with the diphosphate group of the nucleotide sugar, is also conserved in the DUF266 family.

### *SUH1* is the *Arabidopsis* ortholog of the rice *BC10* gene

OsBC10, which has a protein structure most similar to SUH1 (Fig. 5A), is a Golgi-localized type II membrane protein with about 1% of human C2GnT activity in Chinese hamster ovary (CHO) cells (Zhou *et al*., 2009). Before conducting functional analyses, we confirmed that the introduction of the *SUH1-GFP* construct under the *SUH1* promoter restored the phenotype of *ctl1^hot2-1^ suh1-4* plants to that of *ctl1^hot2-1^* mutants (data not shown). This result suggests that SUH1-GFP retains the biological functions of SUH1. To determine the subcellular localization of SUH1, the SUH1-GFP fusion protein was transiently co-expressed with MAN49-mCherry or mCherry-HDEL in tobacco leaf epidermal cells. SUH1-GFP displayed a punctate localization pattern (Fig. 5B), which exactly overlapped with that of the known Golgi marker, MAN49-mCherry, but not with that of mCherry-HDEL localized in the endoplasmic reticulum lumen (Nelson *et al*., 2007). These findings suggest that *SUH1*, similar to the rice *OsBC10*, encodes a Golgi-localized protein containing a DUF266 domain.

**Figure 5.**
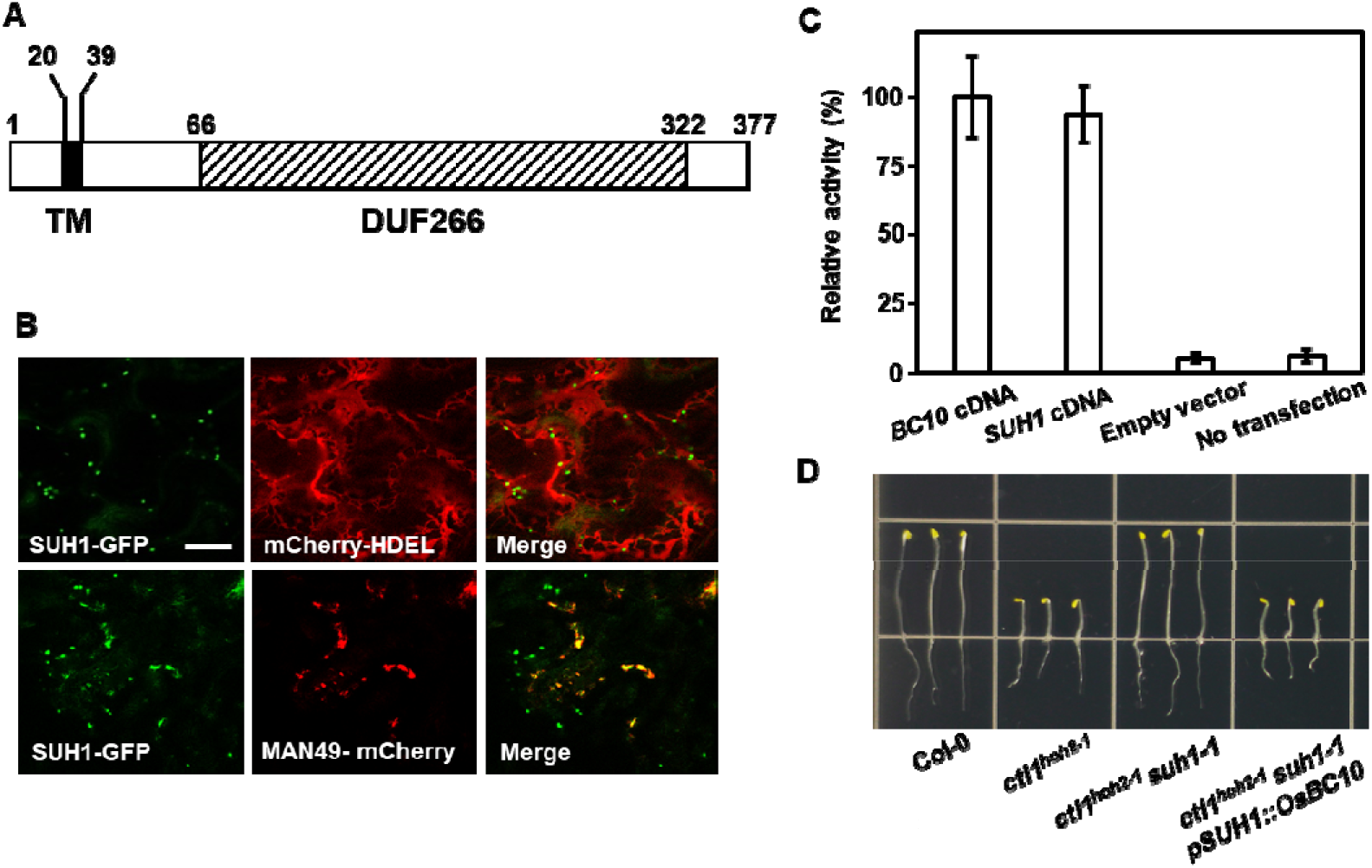
*SUH1* is the *Arabidopsis* ortholog of the rice *BC10* gene encoding a Golgi-localized type II glycosyltransferase. (A) The deduced protein domain structure of SUH1. TM (black box) and DUF266 (hatched box) indicate transmembrane domain and domain of unknown function 266, respectively. The numbers denote the positions of amino acid residues. (B) Confocal images of tobacco leaf epidermal cells transiently co-expressing SUH1-GFP together with mCherry–HDEL or MAN49-mCherry. (C) C2GnT assay in CHO cells. C2GnT activity was determined using PNP-oligosaccharide as an acceptor. The C2GnT activities of each construct and controls were expressed as a percentage of that of BC10. Data are expressed as the mean ± SE of three independent measurements. (D) Complementation of the short-hypocotyl phenotype of dark-grown *ctl1^hot2-1^ suh1-1* mutants with rice *BC10* driven by the *SUH1* promoter. Five-day-old dark-grown seedlings of Col-0, *ctl1^hot2-1^*, and *ctl1^hot2-1^ suh1-1* plants, as well as *ctl1^hot2-1^ suh1-1* plants with *pSUH1::BC10*.

Next, we performed the C2GnT assay using CHO cells as described previously (Bierhuizen and FuKuda,1992). The plasmid carrying either *BC10-GFP* or *SUH1-GFP* construct was transfected into CHO cells. As shown in Figure 5C, CHO cells expressing *SUH1-GFP* displayed C2GnT activity at levels comparable to those transfected with *BC10-GFP*. However, the C2GnT activity was barely detectable in non-transfected CHO cells or in CHO cells transfected with an empty vector. To further assess the potential of *SUH1* as an ortholog of *OsBC10*, *BC10* cDNA under the control of the *SUH1* promoter (*pSUH1::BC10*) was introduced into *ctl1^hot2-1^ suh1-4* mutant plants (Supplementary Fig. S1B). As anticipated, the growth phenotype of *ctl1^hot2-1^ suh1-4* plants expressing *OsBC10* was indistinguishable from that of *ctl1^hot2-1^* mutant plants under both dark (Fig. 5D) and light conditions (Supplementary Fig. S1C). These findings demonstrate that *SUH1*, an *Arabidopsis* ortholog of *OsBC10*, encodes a Golgi-localized type II membrane protein with glycosyltransferase activity.

### *SUH1* is predominantly expressed in tissues associated with the deposition of the secondary cell wall

To investigate the expression pattern of *SUH1*, we generated transgenic wild-type and *ctl1^hot2-1^* mutant plants expressing the β-glucuronidase (GUS) reporter gene driven by the 1,060-bp promoter region of *SUH1* located upstream of the ATG start codon (*pSUH1::GUS*). In the wild-type plants carrying the *pSUH1::GUS* construct, GUS activity was barely detectable in embryos, 5-day-old dark-grown seedlings, and 10-day-old light-grown seedlings with minimal deposition of the secondary cell wall (Fig. 6A-C). However, a strong GUS activity was detected in the interfascicular fibers, xylem cells of inflorescence stems (Fig. 6D), and mature anthers (Fig. 6E), where the secondary cell walls have been reported to be deposited (Taylor-Teeples *et al*., 2015). Similar to the wild-type plants, *ctl1^hot2-1^* showed little *pSUH1*-driven GUS activity in embryos (Fig. 6F) and 10-day-old light-grown seedlings (Fig. 6G), where lignin is not deposited. However, in contrast to the wild-type plants, *ctl1^hot2-1^* exhibited strong GUS staining in etiolated seedlings (Fig. 6H) and pith cells (Fig. 6I), where ectopic lignin is deposited due to the *ctl1^hot2-1^* mutation. An increase in GUS staining due to *ctl1^hot2-1^* was also found in flowers (Fig. 6E, J). These results suggest that the *SUH1* expression is associated with tissues that deposit the secondary cell wall and is enhanced by alterations in cell wall composition caused by the *ctl1^hot2-1^* mutation.

**Figure 6.**
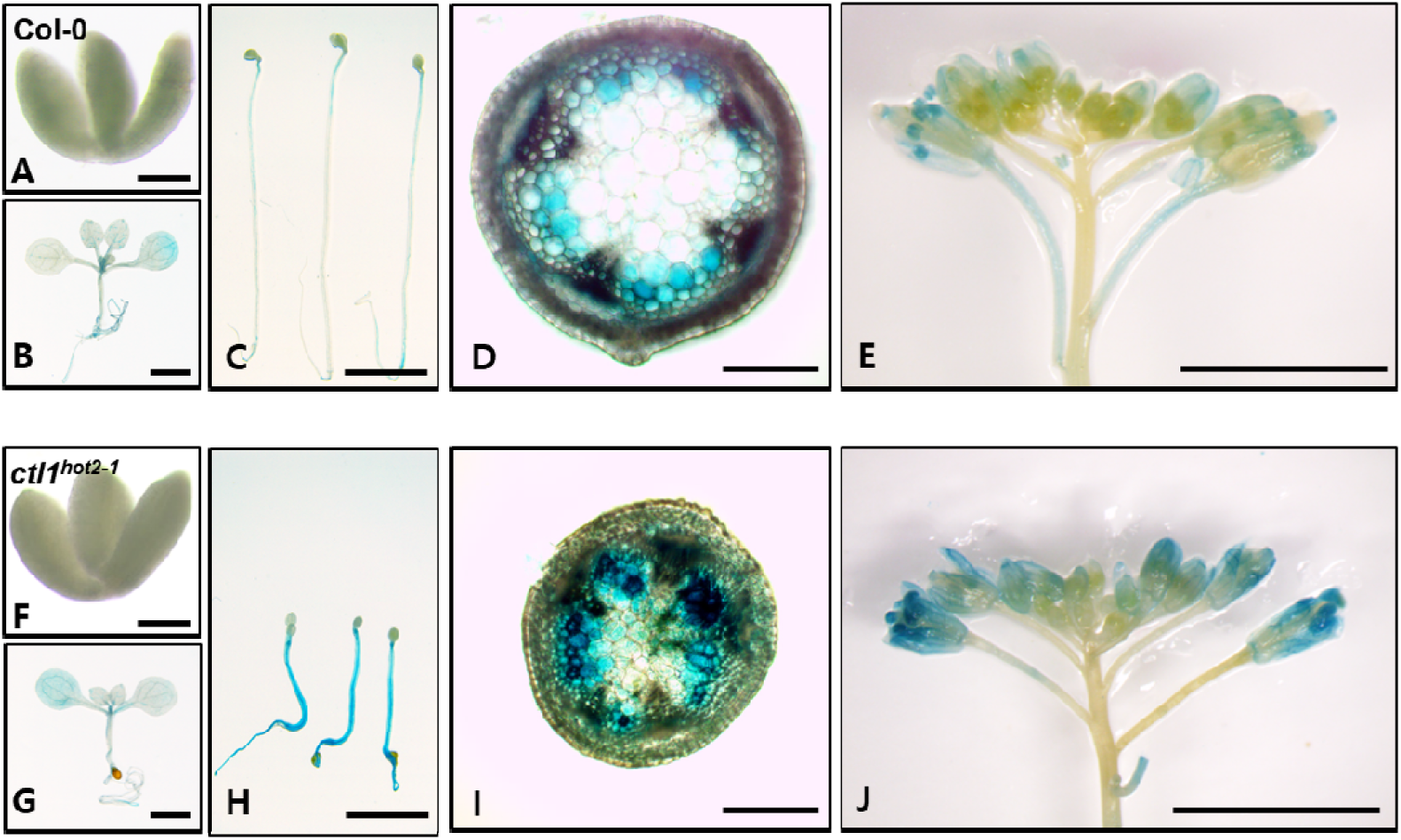
Histochemical assay for GUS activity in transgenic plants with the *pSUH1::GUS* construct. Representative images of Col-0 (A–E) and *ctl1^hot2-1^* (F–G) plants from 15 lines with three replicates. (A, F) The embryos from mature seeds. (B, G) Light-grown 10-day-old seedlings. (C, H) Dark-grown 5-day-old seedlings. (D, I) Hand-cut sections of inflorescence stems of 6-week-old plants. (E, J) Flowers of 6-week-old plants. Scale bars, 200 µm (A, B, C, F, G, and H) and 5 mm (D, E, I and J).

## Discussion

Mutations in *CTL1* cause a number of defects, including growth retardation and changes in cell wall composition in *Arabidopsis*, due to reduced cellulose contents (Hauser *et al*., 1995; Zhong *et al*., 2002; Hong *et al*., 2003). Our findings demonstrate that mutations in *SUH1*, which encodes a Golgi-localized type II membrane protein with glycosyltransferase activity, can restore multiple defects in *ctl1^hot2-1^*mutant plants. These observations provide new insights into the mechanism of cell wall synthesis mediated by *CTL1* in *Arabidopsis*. First, we showed that *suh1*, which has little effect on growth (Fig. 2A and supplementary Fig. S2), causes a non-negligible reduction in cellulose contents even under normal conditions (Fig. 2C) and considerable inhibition of root growth under CBI treatments (Fig. 3). These results suggest that *suh1* also affects cellulose synthesis in *Arabidopsis*, although to a lesser extent than *ctl1^hot2-1^*. If a suppressor mutation displays a phenotype similar to that of the original mutation, the two gene products likely mediate the same step in a multistep pathway (Prelich, 1999). Therefore, considering the genetic relationship between *CTL1* and *SUH1*, it is probable that these two genes are involved in the same process in a multistep pathway.

SUH1 was delivered to the Golgi complex like BC10 of rice (Fig. 5B), while CTL1 was found in various organelles on the secretory pathway from the Golgi complex to the cell wall in *Arabidopsis* (Sánchez-Rodríguez *et al*., 2012). Therefore, the assumption that CTL1 and SUH1 mediate the same step in a multistep pathway suggest the possibility that CTL1-mediated processing takes place in the Golgi complex, where the two proteins coexist, rather than in the apoplastic region where CTL1 is delivered alone. However, our results cannot rule out a possible role of apoplastic CTL1 suggested in a previous report (Sánchez-Rodríguez *et al*., 2012). Interestingly, mutations in *BC15*/*OsCTL1* of rice, the closest ortholog of *AtCTL1*, affect cell wall synthesis by reducing cellulose synthesis, and its product is also targeted to the Golgi complex (Wu *et al*., 2012). These results also make it very interesting to investigate whether the same genetic relationship between *ctl1^hot2-1^* and *suh1* in *Arabidopsis* would also be observed between *bc10* and *bc15* in rice.

Elucidating the substrates of CTL1 is important for understanding its molecular mechanism. Assuming that CTL1 and SUH1 mediate the same step, we consider understanding the possible substrates of SUH1 to be equally important. We showed that SUH1 contains DUF266, which is known to be related to the GT14 family (Fig. 4B), and exhibits C2GnT activity (Fig. 5C). In animal cells, C2GnT is implicated in the elongation of the core 2 *O*-glycan branch of mucin, analogous to AGP glycans in plants (Cheng and Radhakrishnan, 2011; Basu *et al*., 2013). The side chain of AGP glycans is also documented to be elongated by members of the GT14 family in *Arabidopsis* (Knoch *et al*., 2013; Dilokpimol and Geshi, 2014). Therefore, we suggest the possibility that SUH1 and CTL1 collaboratively mediate the glycosylation of AGPs in *Arabidopsis*, probably by promoting and blocking/degrading the same glycans of AGPs, respectively. We also speculate that the *CTL1* mutation disables CTL1 activity, resulting in excessive glycosylation by SUH1, which affects cell wall assembly. On the other hand, in the *ctl1^hot2-1^ suh1-1* mutant, neither CTL1 or SUH1 functions, leading to wild-type growth due to no excessive glycosylation. Therefore, a proper balance between their opposite activities for glycosylation in the Golgi complex may play an essential role in cell wall assembly in *Arabidopsis*.

Finally, while *AtCTL1* expression is relatively strong in almost all tissues (Hossain *et al*., 2010), *SUH1* is predominantly expressed in interfascular fibers and xylems with secondary cell walls (Fig. 6A-E). In particular, *ctl1^hot2-1^* causes increased expression of *SUH1* along with lignin accumulation in etiolated hypocotyl (Fig. 6H) and stem pith cells (Fig. 6I), which are not observed in wild-type plants (Fig. 6C, D). These findings suggest that *SUH1* transcripts are up-regulated by aberrant deposition of cell wall components caused by *ctl1^hot2-1^*. Therefore, in order to further understand the roles of *CTL1* in cell wall synthesis, it is essential to identify its endogenous substrates and elucidate how cell wall integrity affects the expression of *SUH1*.

## Author contributions

Design and project management: SWH

Experiments and data analyses: NTT, HJK, SWH

Writing and editing: NTT, HJK, SWH

## Abbreviations

AGP: arabinogalactan protein
Arg: arginine
*BC10*: *brittle culm 10*
*BC15*: *brittle culm 15*
C2GnT: core 2 β 1,6-N-acetylglucosaminyltransferase
CBI: cellulose biosynthesis inhibitor
*cesa6*: *cellulose synthase6*
CHO: chinese hamster ovary
CTL: chitinase-like protein
DCB: 2,6-dichlorobenzonitrile
DUF266: Domain of Unknown Function 266
GFP: green fluorescent protein
GH: glycosidic hydrolase
Gln: glutamine
Glu: glutamic acid
GT: glycosyltransferase
*irx1*: *irregular xylem1*
GUS: β-glucuronidase
ISX: isoxaben
*ixr1*: *isoxaben-resistant1*
Lys: lysine
prc1: *procuste1*
*qua1*: *quasimodo1*
*SUH*: *suppressor of hot2-1*
*THE1*: *THESEUS1*
Trp: tryptophan

## ACKNOWLEDMENTS.

We thank ABRC for providing the T-DNA insertional mutant for *SUH1* (*At5g14550*, TAIR accession no. SAIL_912_D02) and the seeds of *prc1-1* (Col-O). We also extend our thanks to June M. Kwak (Daegu Gyeongbuk Institute of Science and Technology, Korea) for providing the ER marker gene mCherry-HDEL and the Golgi marker gene MAN49-mCherry. This work was supported by a grant (NRF-2022R1A2C1002724) from the National Research Foundation of Korea.

## Data and materials availability

All data are available in the main text or the supplementary materials.

